# Rhythmic Modulation of Visual Discrimination is Dependent on Individuals’ Spontaneous Motor Tempo

**DOI:** 10.1101/2022.09.10.506584

**Authors:** Leah Snapiri, Yael Kaplan, Nir Shalev, Ayelet N. Landau

## Abstract

Rhythmic structure in our daily experience originates from various sources. It is generated endogenously and observed in spontaneous fluctuations in behaviour and performance. It can also arise exogenously from everyday stimuli, such as speech, motion and music. Here we examined how individual differences in spontaneous motor rhythms affect the tendency to use external rhythmic structure to guide perception. To measure individual differences in spontaneous rhythms of performance we utilized a spontaneous tapping task. To measure individual differences in perceptual rhythmic modulation we designed a visual discrimination task in which targets can appear either in-phase or out-of-phase with a preceding rhythmic stream of visual stimuli. We manipulated the tempo of the visual stream over different experimental blocks (0.77 Hz, 1.4 Hz, 2 Hz). We found that visual rhythmic stimulation modulates discrimination performance. The modulation was dependent on the tempo of stimulation, with maximal perceptual benefits for the slowest tempo of stimulation (0.77 Hz). Most importantly, the strength of modulation was also affected by individuals’ spontaneous motor tempo. Specifically, individuals with slower spontaneous tempi showed greater rhythmic modulation compared to individuals with faster spontaneous tempi. This discovery suggests that different tempi affect the cognitive system with varying levels of efficiency, and that self-generated rhythms impact our ability to utilize rhythmic structure in the environment for guiding perception and performance.

## 1. Introduction

Rhythmic structure is present in everyday stimuli such as speech (Inbar et al., 2020; Poeppel & Assaneo, 2020), motion (Fraisse, 1982; Lakatos et al., 2019), and music (Jones, 2010). These external rhythms can serve as natural cues for the guidance of behavior (Haegens & Zion Golumbic, 2018; Jones, 2010; Large & Jones, 1999; Nobre & van Ede, 2017; Shalev et al., 2019). Numerous studies have demonstrated perceptual benefits for events that appear in synchrony with a preceding beat (Bauer et al., 2021; Cravo et al., 2013; Haegens & Zion Golumbic, 2018; Jones, 2010; Mathewson et al., 2010; Spaak et al., 2014). This phenomenon has been termed “rhythmic facilitation” and was documented in different modalities (Haegens & Zion Golumbic, 2018). However, recent findings emphasize substantial individual differences in rhythm-based perceptual modulation (Bauer, Jaeger, et al., 2015; Doelling & Poeppel, 2015; Lin et al., 2021; Saberi & Hickok, 2021; Sun et al., 2021) . For example, in a study by Bauer and colleagues (2015) only 40 out of 140 individuals showed classical rhythmic facilitation; namely, they performed better for events appearing in-phase with a preceding auditory rhythmic stream, compared to events that appeared out-of-phase (both early and late targets). Similarly, Sun and colleagues (2021) showed rhythmic modulation of auditory detection in 36% of their experimental sample, with no rhythmic facilitation of performance at the group level. Here we aim to test the hypothesis that these individual differences in rhythm-based perceptual modulation are linked to individual differences in spontaneous rhythmic preferences.

Rhythmic patterns in brain and behaviour emerge even when there is no temporal structure in the environment. In the brain, spontaneous rhythmic fluctuations in neural excitability are demonstrated in different timescales from infra-slow to extremely fast (Buzsáki & Wang, 2012; Monto et al., 2008). In behaviour, spontaneous rhythmic structure is found in self-produced motion and speech (Fraisse, 1982; Poeppel & Assaneo, 2020), exploratory behaviours (Amit et al., 2017; Berg, 2002; Moore et al., 2014; Otero-Millan et al., 2008), and cognitive performance (Landau & Fries, 2012; Monto et al., 2008; VanRullen, 2016). Previous work has shown that individual differences in spontaneous rhythmic preferences account for variability in externally paced motor performance (McAuley et al., 2006; Zamm et al., 2015, 2016). For example, McAulay and colleagues (2006) showed that when individuals perform with external tempi that are close to their own spontaneous motor tempo they do better. Namely, they are more accurate and stable in tracking the external rhythmic stimulation. Similarly, in a series of studies Zamm and colleagues showed that musicians perform optimally, both individually, and in a dyad, at tempi that are close to their spontaneous rhythmic preferences (Zamm et al., 2015, 2016, 2018). Interestingly, spontaneous fluctuations in the motor system have been implicated not only in overt-beat tracking, but also in covert beat perception (Cannon & Patel, 2021; Criscuolo et al., 2022). Therefore, individual differences in spontaneous motor tempi might impact not only motor performance, but also modulation of perception within a rhythmic context.

To address this hypothesis, we first assessed individuals’ motor rhythmic preferences using the spontaneous tapping task (Fraisse, 1982; McAuley et al., 2006). Then, we characterized individual differences in rhythm-based perceptual modulation using different tempi of stimulation during a visual discrimination task. Specifically, we designed a visual discrimination task in which targets could appear at different phases with respect to a rhythmic stream of stimuli (i.e., in-phase or out of phase). For each individual we calculated the modulation in discrimination performance as a function of the target phase and the tempo of visual stimulation (0.77 Hz, 1.43 Hz, 2 Hz). Finally, we examined the link between individuals’ spontaneous tapping tempo and the strength of modulation by external rhythmic structure, across the different stimulation tempi.

## 2. Methods

### 2.1. Participants

Fifty-six individuals (55% females, 86.6% right-handed, mean age = 25.2) participated in the experiment. We excluded from the analysis individuals with ADHD (n = 7). No other neurological or psychiatric diseases were reported. In addition, individuals reported normal or corrected-to-normal vision, and normal hearing. We obtained written informed consent from all individuals before the experimental session. Individuals received monetary compensation for their participation. All experimental procedures were approved by the local Ethics Committee of the Hebrew University of Jerusalem.

### 2.2. Experimental tasks

#### Spontaneous tapping task

to assess individuals’ spontaneous rhythmic preferences we used the spontaneous tapping task (Fraisse, 1982; McAuley et al., 2006). We asked participants to tap with their index finger, of the dominant hand, on a smartphone screen, at a comfortable and regular tempo. We recorded tapping times using a touch sensitive app developed in the lab. We placed the smartphone at a comfortable distance on a table positioned in front of the participant. Before the recording we asked the participants to perform a short practice to check that they understood the task. The duration of the recording was 1 minute.

#### Visual discrimination task

to assess the impact of rhythmic context on perception we designed a visual discrimination task. On each trial we presented a rhythmic, isochronous stream of visual stimuli. The last stimulus in each stream was the designated target – an arrow pointing to one out of four possible directions (up, down, left or right). Each stream was comprised of 3-5 preceding events that set the rhythm of stimulation (‘entrainers’). We asked participants to report the direction of the target. We manipulated two aspects of temporal expectation: **(1) Context tempo:** we manipulated the tempo in which the entertainers were flashed on the screen. Overall, we used three tempi of stimulation: 0.77 Hz, 1.43 Hz and 2 Hz, that were presented in three separate experimental blocks. Thus, the inter-onset intervals (IOIs) between subsequent stimuli in a trial were 1.3 sec, 0.7 sec and 0.5 sec for 0.77 Hz, 1.43 Hz and 2 Hz, respectively. The order of experimental blocks was counterbalanced between participants. We selected the tempi of stimulation to correspond with the rhythmic scale of preferred tapping rhythms as established on a separate sample. **(2) Target phase:** we manipulated the timing of the target with respect to the rhythmic context formed by the entrainers. The targets were distributed uniformly across three possible timings: a third of the targets appeared in-phase with the preceding stream (i.e., on beat), a third of the targets appeared half a cycle before the beat (early targets), and a third of the targets appeared half a cycle after the beat (late targets). The different target phases were presented in a random order within each block (i.e., within each context tempo).

### 2.3. Stimuli and experimental protocol

Trial structure and stimulus are depicted in Figure 1. Each trial started with a 0.5 sec fixation dot that was followed by the presentation of a dark-grey diamond (height: 5.5 cm, width: 5.5 cm). The rhythmic stream of stimuli (i.e., entrainers) consisted of light-grey double-sided arrows (length: 5.5 cm, width: 2.8 cm). The double-sided arrows flashed on top of the dark-grey diamond, that was present on the screen throughout the trial. The arrows were presented each for 0.014 sec. The number of stimuli preceding the target stimulus was set to 3, 4 or 5 (10%, 80% and 10% of trials respectively, within each block). We manipulated the number of events before the target to prevent expectations forming based on the number of the stimuli leading to the target. The last stimulus in each sequence was the designated target and was composed of a uni-directional arrow overlaid on the diamond. Participants were instructed to report the direction of the uni-directional arrow immediately after its appearance. When failing to perceive the direction of the arrow participants were asked to guess. Overall, we presented participants with three experimental blocks, each containing a different tempo of stimulation (i.e., context-tempo). Each block consisted of 99 trials uniformly distributed across the three possible target times (in-phase, early, and late targets). Therefore, each participant completed 297 trials. The order of trials within each block was randomized. Breaks were administrated 3 times throughout the task.

**Figure 1.**
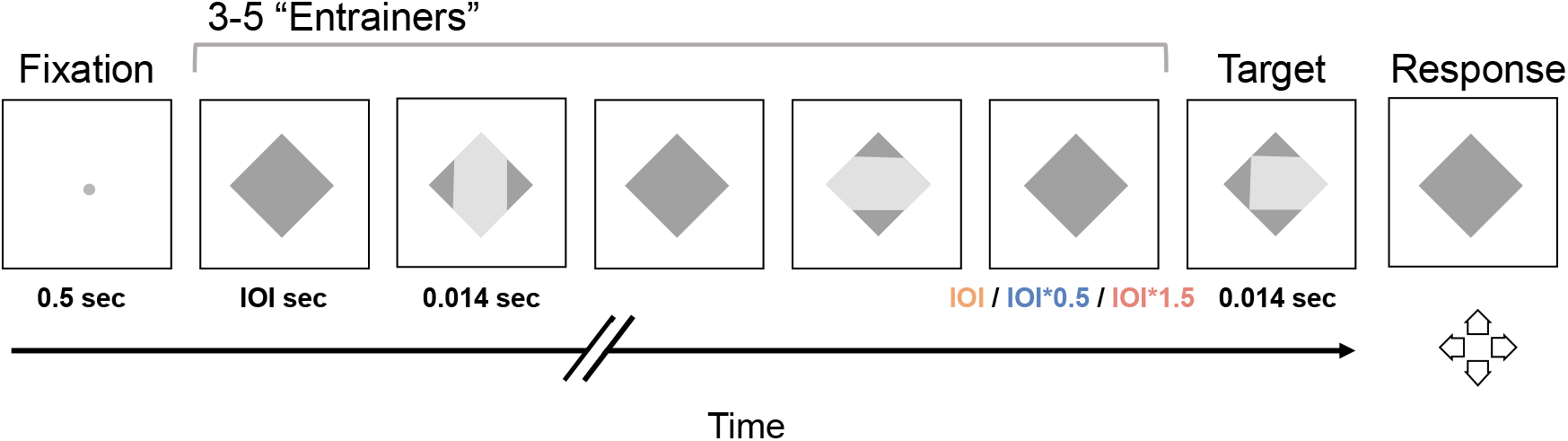
stimuli and trial structure: on each trial, we presented a stream of visual stimuli. The last stimulus in each stream was the designated target – and consisted of an arrow pointing in one direction only. The preceding stimuli consisted of arrows pointing in two directions (i.e., the entrainers). The inter-onset interval (IOI) between the entrainers set the tempo of stimulation and was manipulated across blocks (IOI = 0.5, 0.7 or 1.3 sec). The last IOI before the target was manipulated within each block and could be identical (blue), half a cycle shorter (IOI*0.5, orange), or half a cycle longer (IOI*1.5, pink), than the preceding IOIs in the trial. Before the beginning of each trial, we presented a fixation dot. The entrainers and the target appeared on the screen for 0.014 sec. Participants reported the direction of the target immediately after its presentation.

Participants signed informed consent before arriving to the lab, using a digital form. Upon arrival they completed a personal information questionnaire. Participants were set in a dimly lit room at ∼47 cm from a 60” gamma calibrated monitor and completed the spontaneous tapping task. Then, they continued to the visual discrimination task. Stimulus presentation and response acquisition was controlled using Psychtoolbox (version 3.20.20, (Brainard, 1997; Kleiner et al., 2007) on MATLAB (version 2018b, Mathworks, Natick, Massachusetts). Participants reported the direction of the target using keys that were located at the top of a Logitech joystick (model ATTACK3 simulator). The joystick was located near the participants dominant hand, and participants were instructed to use their thumb for responding. Before starting the task, participants performed a simplified version in which the arrows forming the visual rhythmic stimulation were presented for longer durations (0.100 sec instead of 0.014 sec in the main experiment). This allowed participant to get familiarized with the task and response device. Then, participants continued to a second practice in which the timing parameters were identical to the actual experiment (each stimulus in the stream was presented for 0.014 sec). The stimulation tempo in the practice was always set to the tempo with which the participants will start the experiment. Participants completed ∼ 10 trials of practice before proceeding to the first experimental block.

### 2.4. Data Pre-processing

#### Spontaneous tempo

We calculated the inter-tap intervals (ITIs) between the recorded taps and pre-processed the time course of performance using the following steps: (1) we removed ITIs that were longer than 3 sec, as those were indicative of breaks in performance. (2) we removed ITIs that were more than 1.5 interquartile ranges above the upper quartile or below the lower quartile. On average these procedures resulted in the removal of 2.7% of taps (*SD* = 2.8 %). Then, for each individual we calculated two performance measures: **(1) mean tempo**, calculated based on the inter-tap intervals (ITIs) between the recorded taps. **(2) variability**, calculated based on the coefficient of variation (CV). CV is a normalized measure that accounts for the produced mean tempo in the assessment of variability (ITIs std divided by ITIs mean). Two participants were excluded due to high variability in performance, using the inter-quartile range method for outlier detection. Additionally, we removed two data sets due to a log failure. One of those data sets was replaced by a measurement of the same participant on a different day.

#### Visual discrimination

We removed trials with extremely fast RTs (<150 ms), as those were indicative of premature responses. We did not exclude trials based on other characteristics of RTs, as participants were encouraged to take breaks when needed by withholding the response until they are ready to continue. Overall fast RTs led to the removal of 2.9% of trials. Finally, we removed 2 participants that performed the task at chance level, as was determined using a binomial test. Overall, the analysis was conducted on 44 participants. Analysis conducted on all trials, without excluding premature responses, led to the same results.

### 2.5. Data Analysis

To test the effect of target phase on visual discrimination as a function of context tempo and spontaneous tempo we performed a mixed effect logistic regression (Jaeger, 2008). Our dependent measure was discrimination accuracy, with correct responses coded as 1, and incorrect responses as 0. Our independent measures included ‘target phase’, ‘context tempo’, ‘spontaneous tempo’, and all the interactions between them as fixed effects. It also included random intercepts for participants, and random slopes for context tempo by participant. This random effect structure was the maximal structure to converge (Barr et al., 2013). The two categorical predictors, target phase (before, in phase, after) and context tempo (0.77 Hz, 1.43 Hz, 2 Hz) were both ‘treatment coded’ (reference levels: ‘in phase’ for target phase, ‘1.43 Hz’ for context tempo). The numerical predictor ‘spontaneous tempo’ was centred.

To assess the contribution of each predictor to the model we performed a likelihood ratio test between the maximal converged model and a nested model that excluded the specific predictor we were interested in. In what follows we report for each predictor the Bayesian Information Criteria (BIC) difference, chi-square values and significance level (Meteyard & Davies, 2020).

In addition, to obtain comparisons of interest that were not covered by the contrast structure included in the model, we utilized the ‘emmeans’ package in R (Lenth, 2019). Specifically, we calculated the log-odds difference between specific target phases (before, in-phase, after) for each tempo separately (0.77 Hz, 1.43 Hz, 2 Hz). Then we tested this difference for significance using Wald Z-test and transformed the log odds difference into odds ratio. For each comparison we report odds ratio (OR) with confidence intervals and adjusted significance levels using False Discovery Rate (FDR) correction for multiple comparisons (Benjamini et al., 2001).

Finally, to assess the relationship between individual differences in rhythmic modulation by external rhythms and individuals’ spontaneous tempo we used Pearson correlation. Individual difference in rhythmic modulation were quantified based on the difference in performance between in-phase and out-of-phase targets (separately for early and late targets). This measure emphasizes the impact of rhythmic context, by removing individual differences in baseline performance.

## 3. Results

### 3.1. Rhythmic modulation of visual discrimination is dependent on context tempo

As can be seen in figure 2, the impact of target phase on visual discrimination is highly dependent on the tempo of stimulation. To examine the relationship between context tempo and rhythmic modulation of visual discrimination we performed model comparison between the full converged model and a nested model excluding the interaction term between target phase and context tempo. We found that the interaction contributed significantly to the full model (ΔBIC = 10, χ^2^ (8) = 64.87, *p* <.001). Therefore, context tempo impacts rhythmic modulation of visual discrimination.

**Figure 2.**
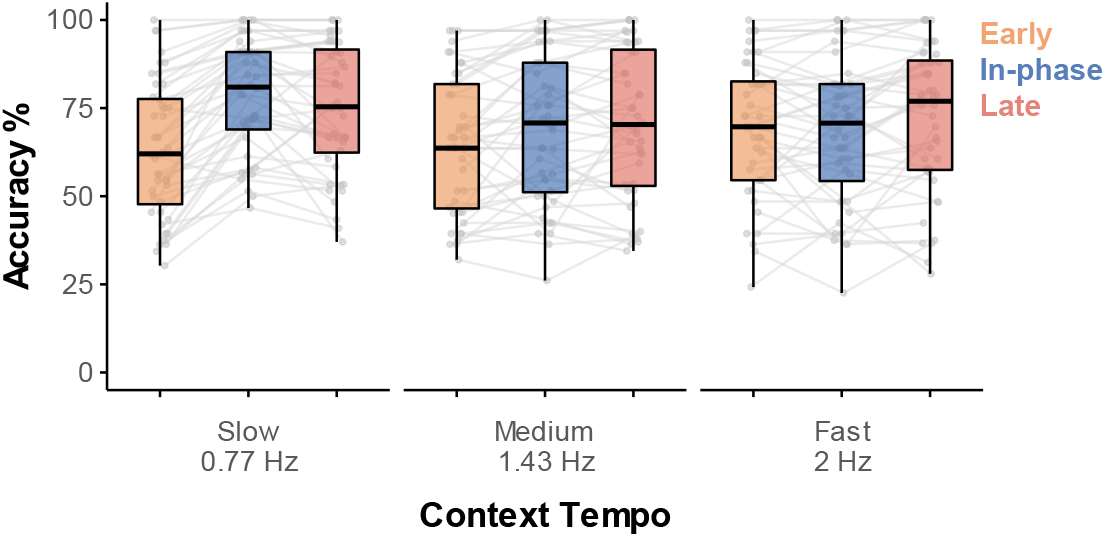
The impact of external rhythms on visual discrimination is dependent on the tempo of stimulation. Participants performed a visual discrimination task. For the slowest presented tempo (0.77 Hz) performance significantly increased for targets appearing ‘in-phase’ with the preceding stream (blue boxplot), compared to targets appearing half a cycle before (orange boxplot), or half as cycle after (pink boxplot). For the medium tempo (1.43 Hz) perceptual benefits were also found for targets appearing in-phase compared to early targets. No perceptual benefits were found for target appearing in-phase with the fast tempo (2 Hz).

To further characterize how performance changes by target phase (early, in -phase, late) for each tempo separately (0.77 Hz, 1.43 Hz, 2 Hz), we obtained log-ratios between groups of interest. We found that the odds to discriminate targets that appear in-phase with the slow tempo (0.7 Hz), are **2.47** times higher than the odds to discriminate early targets (*OR* = 2.47, 95% CI: [2.06, 2.96], *p* <.001), and **1.27** times higher than the odds to discriminate late targets (*OR* = 1.27, 95% CI: [1.05, 1.53], *p* = .02). Therefore, individuals performed significantly better with targets that appeared in-phase with the slow tempo compared to targets that appeared out-of-phase.

Individuals also performed better with targets that appear in-phase with the medium tempo (1.43 Hz), compared to early targets (*OR* = 1.27, 95% CI: [1.07, 1.51], *p* = .01). However, no difference in discrimination performance was found between targets that appeared in-phase with the rhythmic stream and targets that appeared late (*OR* = 0.93, 95% CI: [0.78, 1.12], *p* = .44). Finally, for the fast tempo (2 Hz), we did not find evidence for rhythmic modulation of performance. Individuals discriminated similarly targets that appeared in-phase with a rhythmic stream and early targets (*OR* = .93, 95% CI: [0.78, 1.11], *p* = .44). Furthermore, individuals discriminated targets that appeared in-phase worse than targets that appeared late (*OR* = 0.79, 95% CI: [0.67, 0.95], *p* = .02).

In summary, the impact of rhythmic temporal expectation on visual discrimination is highly dependent on the tempo of stimulation. Maximal rhythmic modulation was found for the slowest tempo we presented with **15.9%** benefit for in-phase targets compared to early targets, and **3.6%** benefit compared to late targets. The medium tempo we presented elicited **4.7%** benefit compared to early targets, and no benefit compared to late targets. The fast tempo did not elicit rhythmic modulation of performance. These results were replicated in another sample with a similar experimental design (Fig 4 in appendix A).

To further assess the impact of context tempo on rhythm-based expectation we compared performance for targets that appeared in-phase with the three different context tempi. We found that the odds to discriminate correctly targets that appeared in-phase with the slow tempo was 1.69 times higher than the odds to discriminate correctly targets that appeared in-phase with the medium tempo (*OR =* 1.69, 95% CI: [1.35,2.12], *p* <.001), and 1.8 times higher compared to the fast tempo (*OR* = 1.8, CI: [1.46,2.24], *p*<.001). Therefore, the slowest tempo we used was the most effective tempo for forming rhythm-based expectation for visual discrimination. We next turn to examine the relationship between rhythmic modulation of performance and individuals’ spontaneous tempo.

### 3.2. Rhythmic modulation of visual discrimination is dependent on individual’s spontaneous tempo

To assess the relationship between individuals’ spontaneous tempo and rhythmic modulation of visual discrimination we performed model comparison between the full converged model and a nested model excluding the interaction term between target phase and spontaneous tempo. We found that spontaneous tempo significantly affected the formation of rhythmic facilitation (*ΔBIC* = 43, χ^2^ (6) = 13.01, *p* =.04). To further breakdown this significant interaction between spontaneous tempo and target phase we calculated Pearson correlation between individuals’ spontaneous motor tempo and the strength of behavioural modulation by target phase. We calculated the strength of behavioural modulation by calculating for each individual the difference in accuracy between targets that appeared in-phase and targets that appeared out-of-phase, averaged across the three different context tempi. This measure emphasized the impact of rhythmic stimulation by removing individual difference in baseline performance. We found that the strength of behavioural modulation is linked to individuals’ spontaneous tempo (Figure 3, *in-phase to early targets*: *r* (42) = .45, *p* = .002, *in-phase to late targets*: *r* (42) = 0.43, *p* = .004). Specifically, individuals with slower spontaneous tempi exhibited greater difference between in-phase and out-of-phase targets, with reduced performance for out-of-phase targets (difference between in phase and out of phase performance greater than 0).

**Figure 3.**
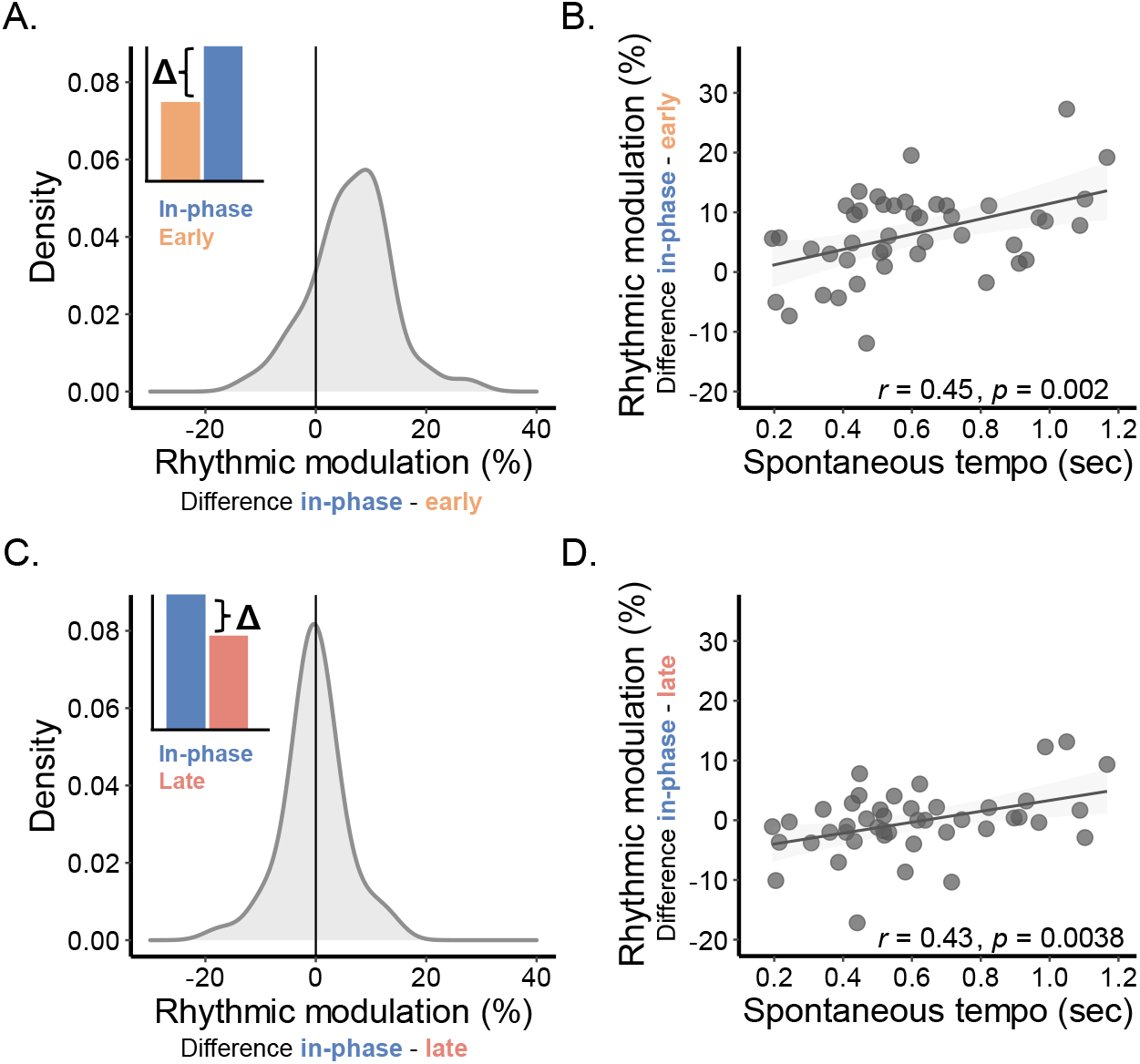
Behavioral modulation by external rhythms is dependent on individuals’ spontaneous tempo. **(A)** Density plot of the rhythmic modulation index, as calculated based on the difference in performance (accuracy scores) between **in-phase** and **early targets**, across all context tempi. **(B)** individuals’ rhythmic modulation index (difference in performance between **in-phase** and **early targets**, y-axis) is dependent on individuals’ spontaneous tempo (assessed through the spontaneous tapping task, x-axis). **(C)** Density plot of the rhythmic modulation index calculated based on the difference between **in-phase** and **late targets**. As can be seen, the group effect is centered at zero (no difference between in-phase and late targets). However, figure **(D)** shows that the direction and magnitude of the rhythmic modulation is dependent on individual’s spontaneous tempo. Therefore, there are substantial individual differences in the impact of external tempo on visual discrimination. This variability can be captured through individual differences in spontaneous motor tempi (i.e., individuals’ spontaneous tempo).

### 3.3. The relationship between spontaneous tempo and rhythmic facilitation is independent of context tempo

We next turned to assess whether the relationship between spontaneous tempo and target phase is dependent on the tempo of external stimulation. To this end we performed model comparison between the full model and a nested model excluding the three-way interaction term between target phase, spontaneous tempo, and context tempo. This interaction did not contribute significantly to the model (*ΔBIC* = 33, χ^2^ (4) = 3.88, *p* =.42). A closer inspection of the correlation between accuracy and spontaneous tempo, for each target phase and context tempo separately indicated a similar relationship between spontaneous tempo and target phase across the different context tempi.

We also tested for the presence of a relationship between context tempo and spontaneous tempo independently of target phase. We compared the full model with and a nested model excluding the interaction between context tempo and spontaneous tempo. This interaction was not significant (*ΔBIC* = 52, χ^2^ (6) = 4.64, *p* =.59). Therefore, individual differences in performance between the different context tempi are not accounted for by individual differences in spontaneous motor tempo.

## 4. Discussion

In this study, we asked whether individuals’ ability to use external rhythmic structure for perception is dependent on context tempo, and on individuals’ spontaneous tempo. We found that visual rhythmic stimulation modulates discrimination performance. The modulation was dependent on the tempo of stimulation, with maximal perceptual benefits for the slowest tempo of stimulation (0.77 Hz). Most importantly, the strength of modulation was also affected by individuals’ spontaneous tempo. Specifically, individuals with slower spontaneous tempi showed greater rhythmic modulation compared to individuals with faster spontaneous tempi. This relationship between spontaneous tempo and rhythmic modulation by external rhythms was not restricted or modulated by the specific tempo of stimulation and was best captured by the difference between ‘in-phase’ (i.e., on beat) and ‘out-of-phase’ performance across the different context tempi.

### 4.1. Tempo specificity in rhythm-based perceptual modulation

Albeit individuals can readily synchronize their motor performance with a wide range of tempi (Repp, 2009), we find a clear advantage for slower tempi in guiding visual perception. These results are consistent with a recent study by Zalta and colleagues (2020) that demonstrated optimal performance in an explicit timing task at a stimulation tempo of 0.7 Hz (i.e., beat-discrimination task). These results were interpreted as evidence in favor of the entrainment framework which predicts optimal performance with external rhythms that are close to endogenous oscillatory activity (Farahbod et al., 2020; Haegens & Zion Golumbic, 2018; Large & Jones, 1999; McAuley et al., 2006; Tavano et al., 2022; Zalta et al., 2020). Such models posit a sampling frequency characteristic of a given system (motor or sensory), that constrain the rhythmic range for optimal percentual benefits. Our findings are consistent with this conceptual framework and go beyond explicit timing judgments, to perceptual discrimination within a rhythmic context.

Rhythmic facilitation in the visual modality has also been demonstrated with substantially faster frequencies (∼10 Hz, De Graaf et al., 2013; Mathewson et al., 2010, 2012; Spaak et al., 2014). This putative discrepancy might reflect two different sources for visual perceptual benefits. Given the short temporal scales, it’s possible that entrainment by a 10 Hz stimulus generates local interactions within sensory cortices while delta frequency stimulation results in the recruitment of top-down temporal anticipation (Fries & Bastos, 2021; Haegens & Zion Golumbic, 2018; Shalev et al., 2019). Future work could attempt characterizing rhythmic perceptual modulation at different temporal scales and address common principles and differences.

### 4.2. Individual differences in rhythm-based perceptual modulation

Several recent studies examining the role of external rhythmic structure in guiding perception show high variability across individuals (Bauer, Jaeger, et al., 2015; Lin et al., 2021; Saberi & Hickok, 2021). Different sources of variability were proposed, such as musical experience (Doelling & Poeppel, 2015) and the strength of neural coupling between frontal and auditory brain areas (Assaneo et al., 2019). Here we show that an important source of variability lies in individuals’ spontaneous motor preferences. This strengthens current models of covert beat perception that emphasize the role of the motor system and its spontaneous dynamics in the sensitivity of the perceptual system to external rhythmic structure (Bauer, Kreutz, et al., 2015; Cannon & Patel, 2021; Criscuolo et al., 2022; Grahn & Rowe, 2009; Morillon & Baillet, 2017; Patel & Iversen, 2014; Ross et al., 2016; Schwartze & Kotz, 2015).

Relatedly, according to preferred period hypothesis (McAuley et al., 2006), when an individual is performing a rhythmic task close to their rhythmic motor preference, benefits on the rhythmic task should be maximal. Our results do not provide direct evidence for this hypothesis. They do, however, highlight that spontaneous tempo impacts cognition. We find that task-tempo and spontaneous tempo impact performance orthogonally: slower tempi generate larger performance benefits compared to faster tempi, and individuals with slower spontaneous tempo are more sensitive to rhythmic stimulation. In other words, our study demonstrates that motor rhythmic preferences can determine the impact of externally presented rhythms on sensory processing and thus affect cognition beyond the motor system.

Finally, our findings also extend current literature on the functional role of spontaneous rhythmic preferences. Previous work showed that spontaneous motor tempi can affect externally paced motor performance in personal (Bardy et al., 2015; McAuley et al., 2006; Roman et al., 2021; Scheurich et al., 2018) and inter-personal settings (Alderisio et al., 2017; Roman et al., 2021; Zamm et al., 2015, 2016, 2018). In addition, previous work investigating the neural markers of rhythm perception found that neural markers modulated by the presence of a rhythm were also linked to individuals’ spontaneous rhythmic preferences (Schwartze & Kotz, 2015). Here, for the first time to our knowledge, we show that individuals’ spontaneous motor tempo impacts visual perception within a rhythmic context, even when overt-beat tracking is not necessary or beneficial for task performance. Therefore, individual differences in spontaneous motor rhythms might be linked to performance variability in a variety of scenarios such as lab-based experiments that include rhythmic temporal structure (e.g. – the statistical learning tasks, see Kirkham et al., 2002), computerized diagnostic tools (e.g., CPT, see Klee & Garfinkel, 1983), and everyday interpersonal interactions that often require rhythmic perception and performance (e.g. communicating, moving and performing together, Keller et al., 2014).

## Data availability statement

The data that support the findings of this study and the code that was used to analyse it will be publicly available after publication on the OSF page https://osf.io/bqu6a/.

## Acknowledgments

This work was supported by funds from Joy Ventures, the James McDonnell Scholar Award for Understanding Human Cognition and the Israeli Science Foundation. The authors would like to thank Yoel Gordon for assistance with data collection, Nitzan Guy, Henry Brice Nir Ofir and Maya Inbar for support in data analysis. We would like to thank Noam Swartz for data visualization advice. We would like to thank Noa Itzhaki for her insightful comments on an early version of the manuscript.

## Author Contributions

LS, SN, and ANL conceived and designed the study. YK collected the data. LS analysed the data. L.S and ANL wrote the manuscript.

