## supplemental figure 4 for "Rhythmic Modulation of Visual Discrimination is Dependent on Individuals’ Spontaneous Motor Tempo"

### Appendix A

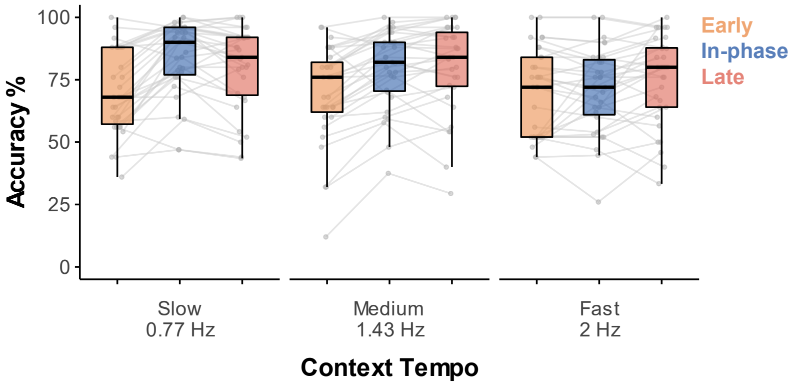


**Figure 4. Replication of the impact of different tempi of stimulation on visual discrimination.** We present here results from an additional sample of thirty-ones participants. In this sample, participant performed the visual discrimination task with two main modifications: (1) the fast tempo was set to 2.5 (compared to 2 Hz in the manuscript above). (2) the distribution of target timings was set to 50% in-phase and 50% out-of-phase targets (25% early and 25% late targets). The pattern of results across the different target phases and tempi are highly consistent with the main report (presented in figure 2 in the manuscript): For the slowest presented tempo (0.77 Hz) performance significantly increased for targets appearing 'in-phase' with the preceding stream (blue boxplot), compared to targets appearing half a cycle before (orange boxplot), or half as cycle after (pink boxplot). For the medium tempo (1.43 Hz) perceptual benefits were also found for targets appearing in-phase compared to early targets. No perceptual benefits were found for targets appearing in-phase with the fast tempo (2 Hz). For detailed description of the experimental design, analysis and results see the following link: https://osf.io/bqu6a/.
